# The activation function of CA3 pyramidal neurons is optimal for the stable recall of memory patterns

**DOI:** 10.64898/2026.09.17.752025

**Authors:** Uri Cohen, Maria Magdalena Picher, Andrea Navas-Olive, Máté Lengyel

## Abstract

Hippocampal area CA3 is widely believed to serve a core memory function: retrieving distributed patterns of neural activity that were previously stored in the recurrent connections between neurons. However, it remains unknown how the physiological properties of individual neurons contribute to this function. Here, we present a mathematical analysis of a canonical recurrent neural network model of CA3. Our analysis provides three main, experimentally testable predictions for recurrent circuits performing stable memory recall. Two of these predictions, that pyramidal cells should be characterized by elevated intrinsic excitability and by inhibition-dominated recurrent connections, are consistent with previous experimental findings. We thus focused on testing the third prediction: that neuronal activation functions have an exponent slightly above 1. For this, we performed *in vivo* intracellular recordings of 49 pyramidal neurons from the CA3 area of behaving mice. Fitting a range of parametric models to these voltage traces and performing statistical model comparison revealed an average neural activation exponent slightly above 1, confirming our theoretical predictions, with substantial heterogeneity across cells, which remains to be accounted for by the theory. An analysis of 133 further cells from two other previous studies provided neural activation exponent estimates highly consistent with those we found in our data. Our results suggest that the properties of hippocampal CA3 area are tuned to support reliable memory recall even at the level of single neurons.

## Introduction

Theory helps us understand complex problems by reaching perfect clarity in simple mathematical models, which are useful if the understanding gained can be extrapolated beyond the model, used to make predictions and explain experimental results in real-world systems. Over the last 50 years, the study of biological associative memory has benefited from challenging the theory to make testable predictions and applying theoretical considerations to understand experimental results.

Based on anatomical considerations, it was suggested that area CA3 of the hippocampus is an auto-associative memory device (1–3). Considerable evidence supports individual components of this hypothesis (4–8), but there is still no unified, quantitatively predictive account connecting single-neuron properties, recurrent synaptic plasticity, and moment-to-moment memory operations.

The seminal work of Hopfield (9) has suggested that binary activity patterns can be stored in the recurrent connectivity of a neural circuit in a way that a partial or noisy memory cue presentation would result – through the dynamics of the network – in a recall of the entire previously stored pattern. This line of work was fruitful in explaining some of the observed properties of cortical and hippocampal circuits, such as the benefit of memory patterns sparseness (i.e., having only a fraction of the population active in each memory pattern (10)), as well as the patterns of synaptic connectivity in CA3 (11), where theoretical results match experimental findings (12, 13), but cannot make predictions on single-neuron properties or the statistics of rate-coded memory patterns.

In a recent theoretical analysis, we relate the ability of a recurrent network to perform stable recall of previously stored rate-coded memory patterns to the statistics of these patterns and the properties of the neural activation function(14). Here we apply this theory to understand the circuit and single-neuron properties in area CA3 of the hippocampus. By considering area CA3 as a natural testbed for our theory, we aim to explain previous experimental results and make new testable predictions. We support one of these predictions by analyzing new and existing CA3 data, while previous studies support the other two predictions (see Discussion).

## Results

### Conditions for stable memory recall of graded memories in recurrent networks

We first summarize relevant results from our previous study (14). Our theory outlines the conditions required for recall of graded (i.e., rate-coded) activity patterns using simple recurrent network dynamics, rather than the previously analyzed binary patterns. We assume the neural activations in each pattern are sampled independently from a log-normal distribution (in line with experimental results (16) from different hippocampal regions and cell types), with a fixed mean (1 Hz) and coefficient of variation, denoted CV (Fig. 1A). Each pattern is sparse; denoting the sparseness level *f*, a fraction 1 − *f* of the *N* neurons in each pattern are exactly zero (arrow at 0 in Fig. 1A and Fig. 1B), and a fraction *f* of the neurons have positive, log-normal activations (Fig. 1A).

**Fig. 1.**
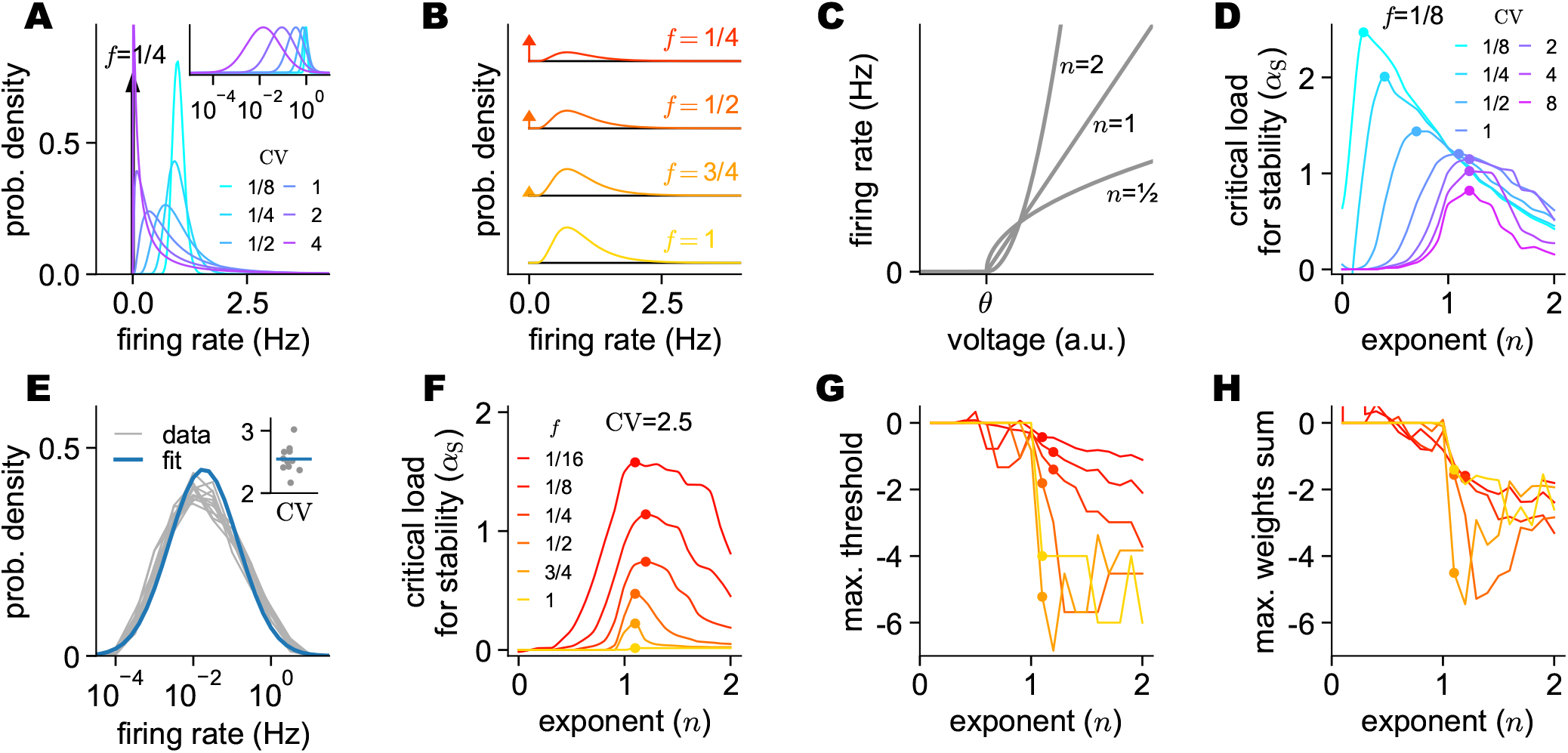
Theoretical results. (A) Probability densities for log-normal distributed firing rates of memory activation patterns, with different CV values (color-coded), and sparseness *f* (arrow at zero firing rate, value indicated). Inset: normalized densities for non-zero firing rates on a log scale. (B) Probability densities for log-normal distributed firing rates of memory activation patterns, with different sparseness values (arrow at zero firing rate). (C) Parametrization of the activation function per Eq. 1, with a threshold *θ* and different choices of the exponent *n* (value indicated). (D) Simulation results for the critical load for stability, *α*_S_, at different choices of the exponent *n* (x-axis) and pattern CV values (color-coded). Using a fixed sparseness *f* (value indicated) and the optimal threshold *θ*. The maximal value in each curve is indicated by a dot. (E) Analysis of neural data from *Alme et al*., *2014* (15). Probability densities for the firing-rate distribution of putative pyramidal neurons from area CA3 in 11 rooms (gray curves), and a fit of a single log-normal distribution to all rooms. Inset: the measured CV in the 11 rooms. (F) Simulation results for the critical load for stability, *α*_S_, at different choices of the exponent *n* (x-axis) and sparseness *f* values (color-coded). Using a fixed pattern CV (value indicated) and the optimal threshold *θ*. The maximal value in each curve is indicated by a dot. (G) The maximal value of the set of thresholds *θ* achieving the values indicated in (F), using the same color-code. The value which corresponds to the maximal critical load in each curve is indicated by a dot. (H) The maximal value of the set of scaled mean weights *Nf* ⟨*W* ⟩ achieving the values indicated in (F), using the same color-code. The value which corresponds to the maximal critical load in each curve is indicated by a dot.

The memory circuit is a general class of recurrent networks of *N* neurons with either voltage or rate dynamics, as those two forms are mathematically equivalent in our case (17). The single-neuron activation function is parametrized by a threshold *θ* and an exponent *n* (Fig. 1C):

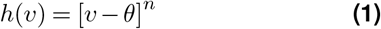

where [*x*] = max{*x*, 0} denotes rectification, and we assume homogeneous parameters across all neurons.

In this setting, we can embed many activity patterns into the network’s recurrent connectivity weights so that those patterns are fixed points of the dynamics, and consider the emergent dynamical stability of these fixed points. Dynamical stability is a technical term for “pattern completion” in memory research: when an activation pattern is a stable fixed point, a partial or noisy cue that resembles the stored pattern would activate the entire pattern. We found a “critical load for stability”, which we denote *α*_S_ and which depends on the above parameters (sparseness *f*, patterns CV, exponent *n*, and threshold *θ*). When the number of patterns *P* is smaller than *α*_S_*N*, these patterns are likely to be stable, and when the number of patterns exceeds this limit, those patterns are likely to be unstable, so a recurrent circuit optimized to be an auto-associative memory would have large *α*_S_ (14).

### Prediction of the pyramidal neurons activation function in area CA3

Going beyond previous results (14), we can now use this theory and the ability to measure *α*_S_ in simulations in order to make predictions about recurrent circuits in the brain. Our computational results show that the critical load for stability decreases with the pattern CV (Fig. 1D), so increased within-pattern variance reduces the number of memories that can be stored and stably recalled. The emerging view of the optimal exponents is non-trivial: for CV *<* 1 the optimal exponent is sub-linear, *n*\* *<* 1. Therefore, when patterns become binary (i.e., very small CV), the optimal choice for the activation function becomes a step function (i.e., a very small exponent *n*). On the other hand, when CV ≥ 1, the optimal exponent is approximately linear, *n*\* ≈ 1, or slightly larger. Under such interpretation, Hopfield-type models (18) may represent the optimal choice for very low within-pattern variance, while our approach is more general and accounts for the full range of pattern variance.

It is thus natural to ask what are the statistics of memory patterns in area CA3 of the hippocampus, noting that these statistics have implications for the number of memories such a circuit can store and stably recall. It is not known which activity constitutes recall of previously stored patterns, and thus which neurons and which time windows should be included in such an analysis. We therefore assume that the statistics of ongoing hippocampal activity are representative of the statistics of memory patterns. This activity has previously been found to be log-normal distributed (16, 19). To exactly characterize the statistics of ongoing CA3 activity, we have re-analyzed previously collected electrophysiology results from area CA3 in which rats attended 11 different rooms and the activity of *N* = 332 putative pyramidal neurons was recorded extracellularly (15). Assuming that activity in a given room is representative of a putative memory pattern, we find that their population density matches a log-normal distribution (Fig. 1E), in line with previous literature (16, 19) and our assumptions on memory pattern statistics (Fig. 1A inset). Furthermore, the measured coefficient of variation was 2 − 3 in all 11 rooms (Fig. 1E inset). We note that 71-95% of neurons were active in each room, but the sparseness of specific memory patterns is expected to be much higher (i.e., *f* would be smaller), both because electrophysiology is biased toward active neurons and because we measured the sparseness of ongoing activity under unrestricted behavioral conditions and over long experimental periods.

In the theory, we assumed that activity is independent across memory patterns, but the experimental results diverge from this assumption, as single-neuron activity is correlated across rooms. We find cross-room correlations of *c* = 0.35 for the log-firing-rate of different neurons, but not cross-neuron correlations for different rooms (Fig. S1A-C). We thus repeated our simulations with the observed level of cross-pattern correlations and found that the critical load for stability decreases only slightly due to these substantial correlations (Fig. S1D-F). This result is surprising, as in the Hopfield model, for binary patterns, such correlations substantially reduce the number of memories and require modification of the learning rule (20).

Focusing on the biological regime found above, CV = 2.5, our model demonstrates that the critical load for stability increases with pattern sparseness (Fig. 1F), reproducing the well-known benefit of such patterns in the binary patterns case (10, 21), and biological findings of sparse activity of pyramidal neurons in the hippocampus of rodents (22, 23). Furthermore, the optimal exponent is always slightly above 1 (Fig. 1F), and the associated threshold *θ* is always negative for supra-linear exponents *n* ≥ 1 (Fig. 1G), including the optimal one. Similarly, the sum of the synaptic weights to each neuron is always negative for supra-linear exponents *n* ≥ 1 (Fig. 1H), including the optimal one.

The above simulation results are complemented by analytic results for a more restrictive case of dense patterns (14), and both cases make the same predictions for the biology-relevant case of high-variance memory patterns. We make the following three predictions regarding pyramidal neurons in hippocampal area CA3:

1. The activity threshold should be negative, so that neurons should demonstrate “positive excitability”, a spontaneous firing without recurrent inputs.
2. The mean synaptic weights for each neuron should be negative, so in biological terms the synaptic weights are inhibition-dominated.
3. The neural activation function should have an exponent *n* which is 1 or slightly above 1, i.e., close to threshold-linear.

The two former predictions are supported by the literature (see Discussion). We thus focus on the latter prediction and directly test it in what follows.

### Collecting subthreshold voltage from *in vivo* recording in behaving animals

Step-current injections are commonly used *in vitro* to reconstruct a neuron’s activation function (denoted “f-I curve” in this context). However, such analysis falls short in determining the properties of the activation function under *in vivo* conditions, as changes in the variance of the background noise affect the neuron’s activation function (24). The common interpretation of this observation is that, while the intrinsic spike-generation mechanism of cortical neurons is highly reliable, the stochasticity observed *in vivo* is due to fluctuations in the synaptic drive (25). To account for the stochastic, irregular spike times observed *in vivo*, it was suggested that neurons operate in a regime where excitatory inputs are continuously counterbalanced by inhibitory inputs, so the timing of threshold crossings becomes highly stochastic (26, 27). Regardless of the precise mechanism (e.g., deterministic chaos, true randomness), the evidence identifies the recurrent network architectures as the source of noise in biological networks (28).

Characterizing the activation function can be performed *in vivo* using step current injections (e.g., 29) or optogenetic activation (30, 31), but such a protocol is expensive in terms of the number of samples collected per unit time. Instead, we opt for an analysis method that relates a neuron’s subthreshold voltage to its firing, for which the natural fluctuations of a neuron’s voltage are an advantage. Similar methods were previously used to study the response of vision neurons (32, 33). This method requires access to their intracellular voltage *in vivo* while the animals are awake and behaving, and allows for characterizing the activation function of CA3 pyramidal neurons under conditions relevant to memory recall.

We thus performed whole-cell patch-clamp recordings *in vivo*, in awake mice running on a linear belt (Fig. 2A). For intracellular recordings, a patch-pipette was positioned in the hippocampal CA3 subfield, while a silicon probe featuring 32 recording channels spanned all subfields of the hippocampus. The patch-pipette contained biocytin for post-hoc labeling and locating the recorded neuron (Fig. 2B). Recorded cell locations spanned the entire proximal-distal axis of CA3 (Fig. 2C). The animal’s velocity and position were monitored during the experiment (Fig. 2D).

**Fig. 2.**
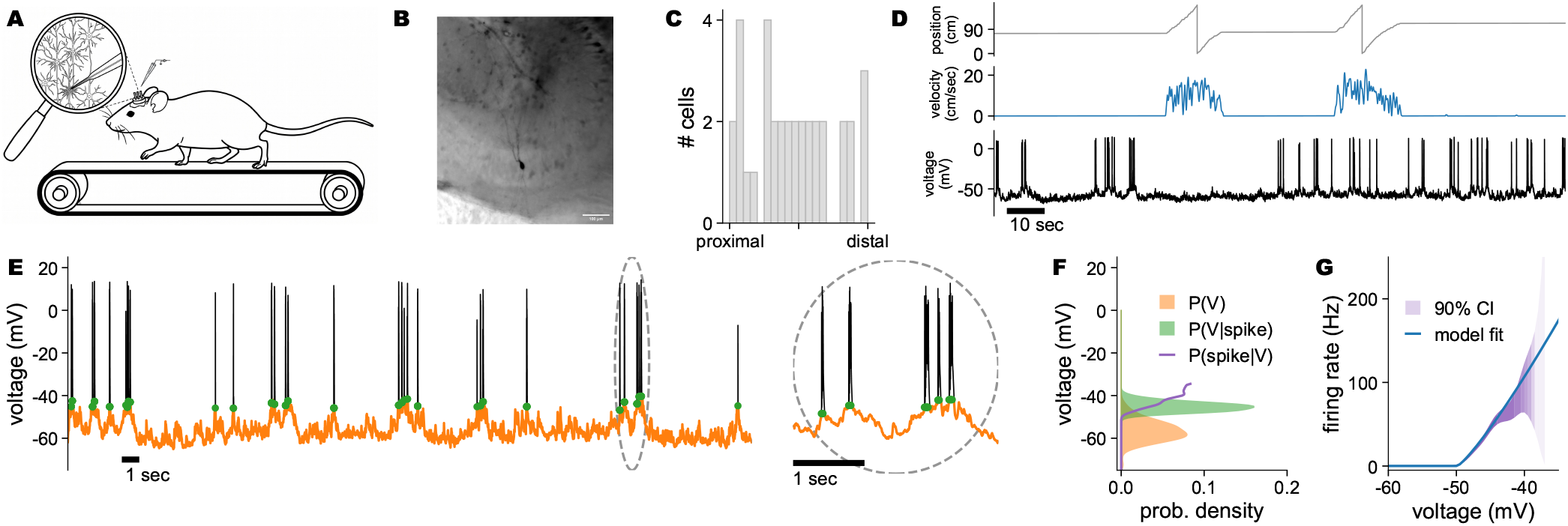
Illustration of the experimental and analysis pipeline. (A) An illustration of the experimental procedure: a mouse running on a linear belt while a patch-pipette is used for whole-cell intracellular recordings from CA3. (B) A photo of a CA3 pyramidal neuron filled with biocytin during recording. (C) A histogram of cell body locations of recorded CA3 pyramidal neurons along the proximal-distal axis. (D) Example traces of position (gray), velocity (blue), and intracellular voltage (black). (E) Example voltage trace, with spikes (black), subthreshold voltage (orange), and voltage at spike initiation (green). Inset: larger view of the circled range. (F) Histograms of the distribution of subthreshold voltage (orange) and the voltage at spike initiation (green). The resulting spike probability for each voltage from Bayes’s rule is presented as overlayed curve (purple). (G) A 90 % credible interval for the firing rate as calculated from the Bayesian analysis (purple; lighter shade indicates a wider interval). The best fit of the parametric model, Eq. 2, assuming inhomogeneous Poisson spiking with instantaneous rate given by the model.

Electrophysiology results from *N* = 49 putative CA3 pyramidal neurons were subsequently processed to analyze the relation between the neuron’s voltage and spiking activity. By identifying precise spike times and separating the subthreshold voltage trace from the spike-related component (Fig. 2E), we could estimate the density of subthreshold voltage *P* (*V*), as well as the exact voltage at spike initiation *P* (*V*|spike) (orange and green curves in Fig. 2F, respectively). From these distributions, using application of Bayes law Eq. 4, the conditional probability *P* (spike|*V*) was extracted (purple curve in Fig. 2F). As the point estimator for the conditional probability is a noisy statistic, we consider the 90 % credible interval around it, which provides an explicit measure of how well it is constrained by the data (narrow or wide purple interval in Fig. 2G).

### Modeling the voltage-to-firing-rate transformation in behaving animals

The conditional probability analysis is non-parametric and does not assume any specific shape for the resulting estimate of firing rate *r*(*V*) = *P* (spike|*V*)*/dt*. By assuming a parametric model:

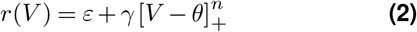

where *ε >* 0 is a noise term to allow for some level of spiking below the threshold, *γ* is a positive gain factor, and the threshold *θ* and exponent *n* are the same as above. While this model is more complicated than the one analyzed in the theory, Eq. 1, we note that both the gain *γ* and noise term *ε* do not change the resulting theoretical predictions (see Methods) and allow us to accommodate some of the biological heterogeneity between neurons.

Assuming the neural spiking follows an inhomogeneous Poisson process with firing rate given by Eq. 2, we can fit the optimal parameters directly from the voltage trace and spike times (blue curve in Fig. 2G). This analysis is applied directly to the voltage and is independent of the non-parametric analysis for the credible interval above. Thus, the agreement between the results of both methods in Fig. 2G is due to the underlying structure of the data.

To demonstrate that this procedure can recover the correct parameters, we generated synthetic spike times from the real subthreshold voltage traces by sampling from an inhomogeneous Poisson process with firing rate given by Eq. 2, using different parameter values. We repeated this procedure multiple times using the entire trace of each cell. The ability to recover those “ground truth” values using our model fit process is demonstrated in Fig. S2.

We applied the two analysis methods to the 49 neurons for which we collected intracellular voltage, and present the per-neuron results in Fig. 3. For each neuron, the credible interval for the conditional probability *P* (spike|*V*) provides a non-parametric estimation of the region in which the activation function may operate, while the model fit further assumes a specific shape of the activation function. We thus use the non-parametric analysis to identify ‘well-constrained’ versus ‘poorly-constrained’ results, based on the width of the credible interval (see Methods). The number of spikes in the recording is the main determinant of how well the results are constrained. To visualize this, we sort the results by the number of spikes per recording and color them by constraint quality.

**Fig. 3.**
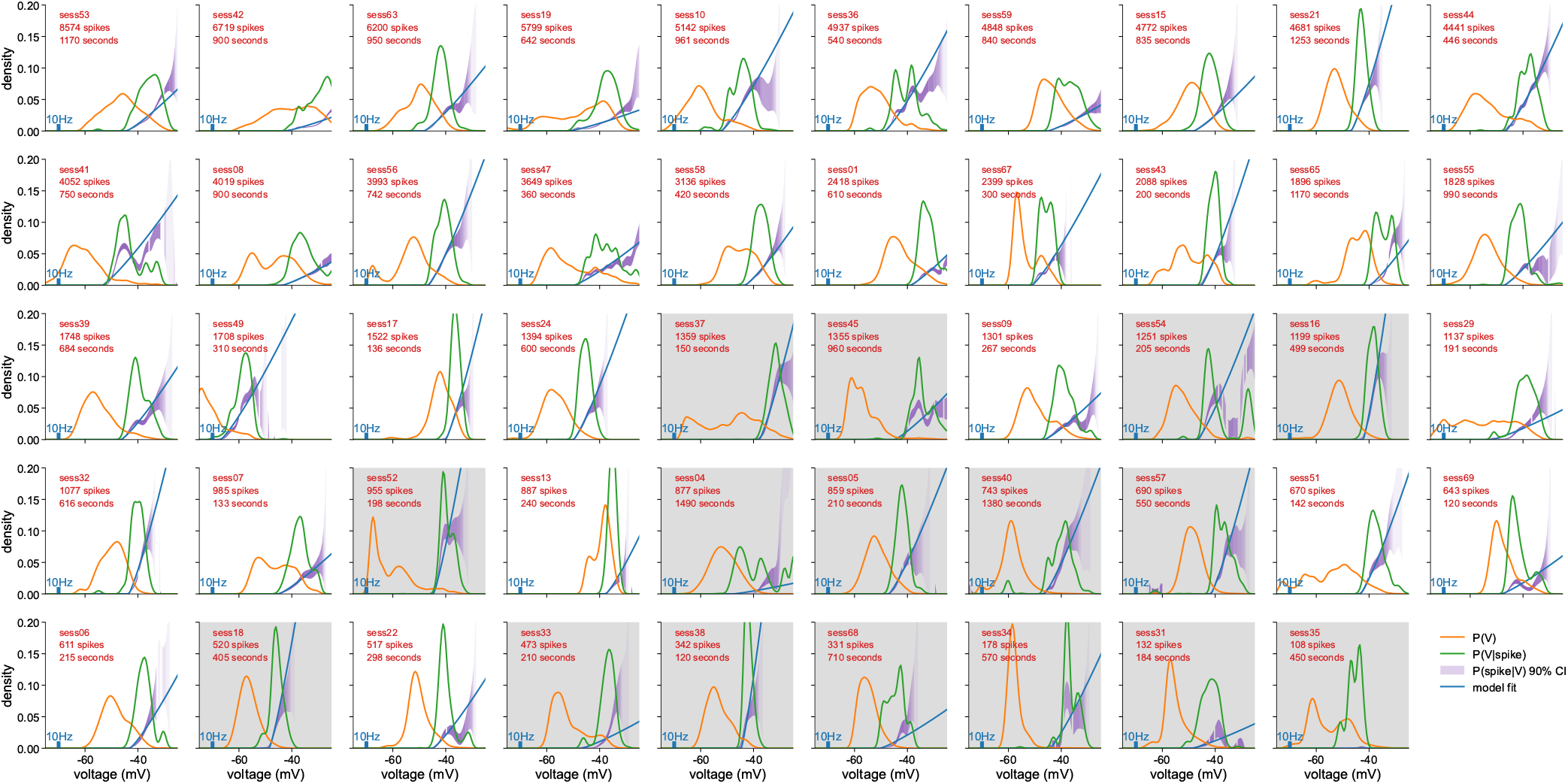
Analysis of voltage-dependent spiking probability in CA3 pyramidal neurons from our dataset. A panel for each intracellularly recorded cell, ordered by the number of spikes. Panel background color is white for ‘well-constrained’ cells, and gray for ‘poorly-constrained’ cells. Each panel presents the distribution of subthreshold voltage (orange), the distribution of voltage at spike initiation (green), and a 90 % credible interval for the resulting spike probability at each voltage (purple; lighter shade indicates a wider interval). The blue curve is the activation function of the best-fit parametric model, Eq. 2, in units of Hz (scale indicated at each panel) or in units of spike probability at each voltage (regular y-axis). The best-fit activation function has *n*\* = 1.19 and *ε*\* = 0.10 shared across all neurons; *θ*\* and *γ*\* were fit for each neuron.

### CA3 neurons’ activation function is best explained as threshold-linear

To maximize the statistical power of our analysis, we curated existing voltage-recording datasets from CA3 in behaving mice (see Discussion). We then applied the same analysis as above to those neurons: *N* = 37 neurons from *Malezieux et al*., *2020* (29, presented in Fig. S3) and *N* = 96 neurons from *Li et al*., *2024* (8, presented in Fig. S4).

We note that the cells in the three datasets (ours, *Malezieux et al*., *2020*, and *Li et al*., *2024*) differ in the number of spikes collected per neuron, due to both differences in the duration of the recordings (Fig. 4A) and differences in the average firing rates (Fig. 4B). Those differences may be related to behavioral procedures; the fraction of the session in which the mouse is in a ‘running’ state (compared with a ‘still’ state, see Methods) differs substantially between the datasets (Fig. 4C), as well as the average velocity (Fig. 4D) and velocity distribution (Fig. S5).

**Fig. 4.**
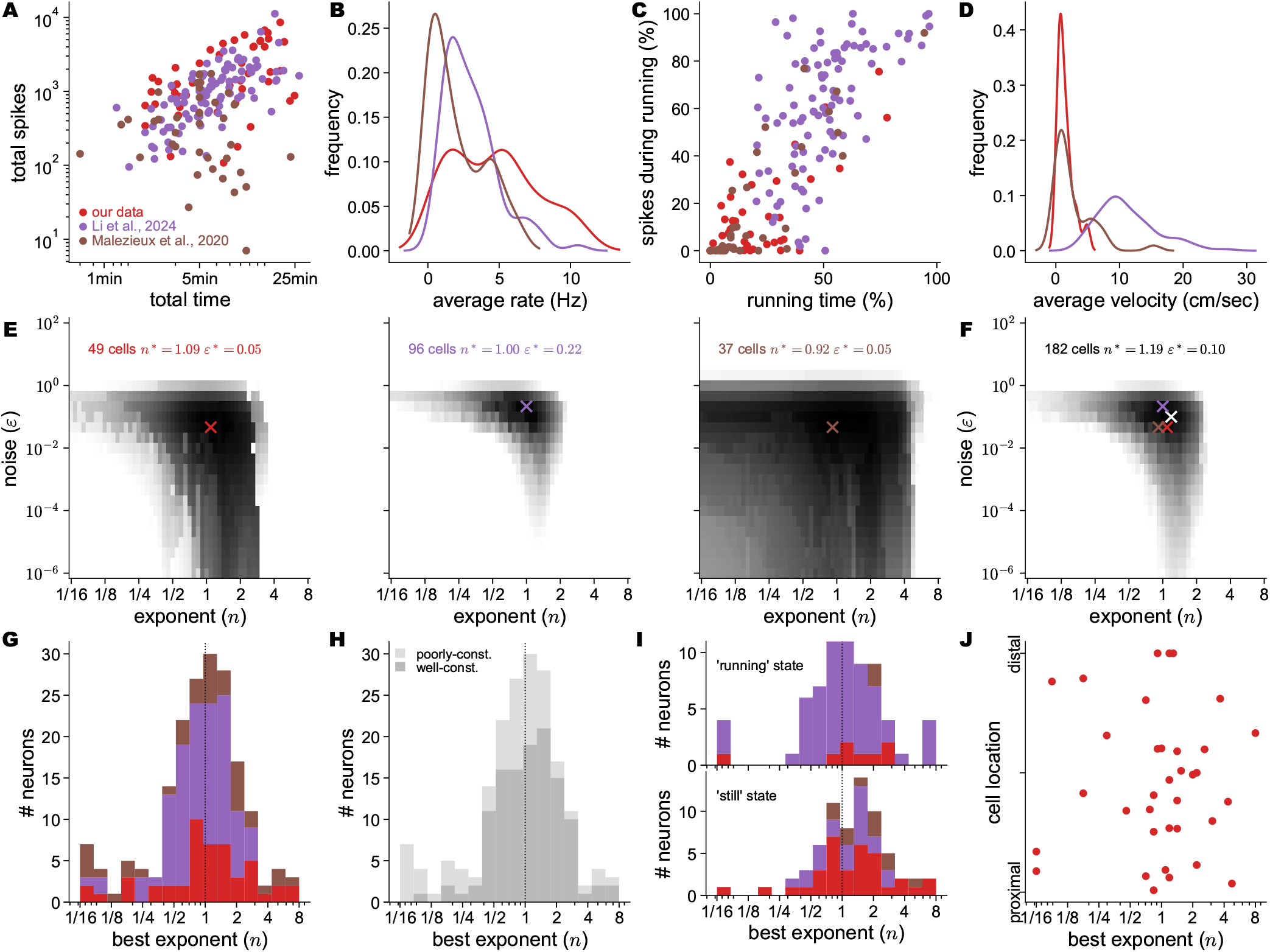
Analysis of optimal exponent fit. (A-D) Comparison of the different datasets, color-coded per the legend in (A). (A) Scatter plot of the total number of spikes versus the total recording time, a dot per session. (B) Histogram of the average firing rate across neurons in different datasets. (C) Scatter plot of the percent of spikes occurring at the ‘running’ state versus the percent of time spent at the ‘running’ state, dot per session. (D) Histogram of the average animal velocity across different datasets. (E-F) Model fit per dataset (Our data, *Malezieux et al*., *2020* (29), and *Li et al*., *2024* (8), (E)) or across all datasets (F) or a model with shared noise *ε* (y-axis) and shared exponent *n* across all neurons (gain *γ* and threshold *θ* are fit per neuron). In each panel, log-likelihood is color-coded in grayscale (white: 25,000 below the maximal value, to black: the maximal value). Crosses (color-coded as in (B)) indicate the maximum likelihood for each dataset (E), and are reproduced in (F), along with the maximum likelihood value across all datasets (white). (G-H) Histograms of the number of neurons versus their optimal per-neuron exponent *n*\*, colored by dataset (G) or by how well-constraint the results are (H). Using a model with private noise *ε* and private exponent *n* (thus all parameters are fit per neuron). (I) Like (G), for a model fitted to voltage from the ‘running’ state (top) or ‘still’ state (bottom), using only ‘well-constraint’ results. (J) Scatter plot of the optimal exponent *n*\* from (G) versus cell locations along the proximal-distal axis of CA3.

For the sake of simplicity, we first consider the case where a single exponent *n* and a single value *ε* are shared across neurons. Here and throughout our analysis, we assume that the gain *γ* and threshold *θ* may be different across neurons. Under these assumptions, Fig. 4E presents the optimal choice for *n* and *ε* in each dataset, and Fig. 4F presents the optimal choice across all neurons. Despite the observed differences between datasets (Fig. 4A-D), we find a considerable similarity in the recovered optimal values (crosses in Fig. 4F).

The resulting optimal values in this analysis are an exponent *n* = 1.19 and a noise term *ε* = 0.1Hz, so that the exponent is in great agreement with the theoretical prediction (Fig. 1F). Interestingly, the optimal value of *n* across all neurons is slightly larger than the optimal value per dataset, presumably due to interaction between the shared noise *ε* and the shared exponent *n*. We have used those fixed values to demonstrate the model fit in Fig. 3, Fig. S3, and Fig. S4. Despite using a single exponent and a single noise term to match the properties of all neurons across three datasets, the credible interval aligns reasonably well with this model fit in these results.

Beyond a single set of shared parameters, we would now consider different statistical models more systematically. The first kind of model assumes a specific value of *n* and fits other parameters per neuron. In agreement with our prediction, under different model-comparison criteria (see Methods), *n* = 1 is favorable compared with both *n* = 1*/*2 and *n* = 2 (Fig. S6A).

To consider a broader set of models, we note that each parameter, the exponent *n* and noise term *ε*, may be private (i.e., allowed to be different for each neuron), or shared across neurons. Among those four models, different model-comparison criteria favor the model where both parameters are private (Fig. S6B).

We thus focus on this model, where the exponent *n* and noise *ε* are allowed to be different for each neuron. We thus present the histogram of the optimally found exponent *n* across the three datasets in Fig. 4G-H. The histogram is centered around the theoretically predicted *n* = 1, but a substantial number of neurons reside over the entire range of *n* ∈ [1*/*2, 2]. This width cannot be explained by the noisy nature of our inference process, as fitting an exponent *n* on synthetic data with the same statistics as the real data (but known ground-truth values for the activation function) yields a much narrower error (Fig. S2B). Thus, these results are consistent with a certain level of heterogeneity in the firing-rate exponent across the population, with most neurons having an exponent near *n* ≈ 1. Some of this heterogeneity is related to the ‘running’ state; repeating the analysis on voltage traces in this state yielded a wider histogram of optimal exponents, compared with the ‘still’ state (Fig. 4I). Finally, the optimal per-neuron exponent cannot be explained by the neuron location in CA3 (Fig. 4J).

## Discussion

### Known properties of CA3 neurons and circuit support our “positive excitability” prediction

A negative threshold defines an intrinsic drive for activity, which in biological terms can stem from either intrinsic cellular properties or a tonic synaptic drive, with the theory silent on the distinction between these two sources. The defining feature of a negative threshold – or a “positive excitability” – is that neurons should be spontaneously active in the absent of recurrent drive. *In vitro* results from guinea pigs demonstrate that CA3 neurons maintain their spontaneous activity in slice preparations, with pyramidal cells still triggering spontaneous discharges after blocking both excitatory and inhibitory synaptic transmission (34). Furthermore, spontaneous CA3 discharge is intrinsic rather than synaptically driven; action potentials recorded intracellularly from CA3 pyramidal cells were initiated in the absence of a detectable synaptic event, in contrast to action potentials generated by nearby inhibitory cells, which were preceded by an EPSP (34). *In vivo* results from anesthetized rats show CA3 pyramidal neurons exhibit spontaneous firing and bursting activity (which was similar for pyramidal neurons from CA1 and distinct from dentate gyrus (DG) granule cells, lacking spontaneous activity). CA3 neurons had higher resting voltage and higher membrane resistance compared with both CA1 and DG (35).

It should be noted that “intrinsic membrane excitability” differs from our “positive excitability”, as the former may result from a change in gain – increased response to external drive – rather than a change in threshold. For example, *in vitro* characterization of the intrinsic membrane excitability properties along the transverse axis of CA3 has reported a decrease in resting potential along this axis (from the distal CA3a to the proximal CA3c), associated with an increase in input resistance so that the proximal parts are more easily excitable by current injections (36), inline with previous studies (35, 37). It is thus unclear whether this gradient corresponds to a changed activity threshold and, in turn, a changed ability to perform memory recall.

The definitive experiment to test the negative threshold / positive excitability prediction would be to observe spontaneous activity of CA3 pyramidal neurons even after recurrent collaterals and local inhibition are suppressed pharmacologically or optically, but such an experiment has not been conducted, to our knowledge.

### Known properties of CA3 neurons and circuit support our “inhibition-dominated recurrent connectivity” prediction

The main simulations we performed here and in our previous study (14) do not respect Dale’s law and thus make no distinction between excitatory and inhibitory neurons. A possible interpretation in biological terms would be that the theory applies to the pyramidal neurons, with the fast activity of the local interneurons ‘absorbed’ into the connectivity weights. Under this interpretation, predicting inhibition-dominated synaptic connections means that activation of recurrent connections would generate a net inhibitory response in adjacent neurons. This is the result of a classic intracellular recordings study from CA3 principal neurons of cats, where stimulation of recurrent collaterals resulted in a net inhibitory response (38). This effect is reproduced in a recent *in vitro* study in mice (36) finding that recurrent collateral stimulation by either electrical or optogenetic means triggers inhibitory responses that are stronger than excitatory responses by a factor of ×3− ×5, with non-monotonic heterogeneity across CA3 subregions (the middle CA3b region exhibiting a smaller factor compared with both CA3a and CA3c).

The definitive experiment to test inhibition-dominated recurrent connections prediction would be an *in vivo* reproduction of the cat results in mice, using the modern toolset of activity perturbations and measurements.

### Available intracellular voltage recordings from CA3 in behaving rodents

Intracellular voltage recordings from CA3 in behaving rodents are not common in the literature. Pioneering whole-cell patch-clamp recordings in behaving rodents allowed for collecting voltage data from hippocampal principal neurons in head-fixed behaving animals (39). While initially focused on hippocampal area CA1 (40– 44), similar recordings were eventually performed also in area CA3 (8, 29, 45). Voltage imaging with a genetically encoded voltage indicator rather than patch-clamp (30, 31, 46, 47) is currently not available for hippocampal area CA3. As noted in the results, we could apply our analysis to *N* = 37 neurons from *Malezieux et al*., *2020* and *N* = 96 neurons from *Li et al*., *2024* (8).

### The shape of activation function in the brain

Analyzing the relation between voltage and firing rate resulted in a threshold-linear ‘f-V curve’ in multiple cortical areas. For example, the analysis of neural responses in V1 used a rectified model with *n* = 1 (48), whereas an attempt to measure the exponent of these responses yielded *n* = 1.2, and thus coincides with our result (49). Similarly, a rectified-linear model (i.e., *n* = 1) was found to describe well the results in frontal cortex persistent activity, where it was related to attractor-like behavior (50). Optogenetic stimulation methods applied *in vivo* allow researchers to go beyond passive characterization by perturbing activity. In V1, it is possible to apply such perturbations at different network states based on visual properties of the stimulus, and the resulting activation function of visual neurons was shown to be supralinear at threshold, linear at intermediate values, then saturating at high values (51). In hippocampal area CA1, an all-optical approach using increasing light intensity was used to characterize the ‘f-I curve’ of multiple cell types. The resulting curves are linear, with different gains across behavioral states (30, 31).

Those results are consistent with our findings at area CA3 and seem at odds with recent results showing that the potential input-output transformation of neurons is very complicated (52). This discrepancy can be reconciled by noting that our analysis does not rule out the case where the transformation from inputs to somatic voltage can still be complicated. Another possibility is that the reported complexity of an isolated neuron is not reflected in the effective input-output transformation under *in vivo* conditions.

### The dynamical regime of memory recall

While memory storage and recall are well-defined stages in our theoretical model, it is unclear whether they correspond to distinct dynamical modes in the hippocampus. It is thus conceivable that our three predictions about pyramidal neurons in area CA3 hold only when the network is in “recall state”. It was proposed that acetylcholine enhances the influence of input connections relative to recurrent excitatory connections and supports synaptic modification, and thus can be related to the storage of new information (53), and further that those might correspond to different phases within the theta rhythm cycle (54). This seems consistent with the reduction of excitability – voltage hyperpolarization and decreased firing – reported for CA3 putative pyramidal neurons during ‘theta’, in contrast to times of ‘large irregular activity’ (29). Interestingly, our analysis of ‘running’ versus ‘still’ state may have some overlap with the distinction between ‘theta’ and ‘large irregular activity’ (29).

### Comparison with other formalisms

The proposed balance between inhibition-dominated recurrent connections and positive excitability is not equivalent to “tightly balanced excitation and inhibition” (27). In E-I balance, each connection is relatively strong, but the net input mean and variance are both independent of the number of inputs, leading to fluctuation-driven dynamics (often corresponding to the asynchronous irregular state). In our model, each connection is small, with the net input having a finite negative value which is balanced by the positive excitability bias. Our regime supports the iceberg effect of excitability, where changing it smoothly changes the population mean firing; in the E-I balance, increasing external drive (which corresponds to positive bias) causes a phase transition in the network’s dynamical regime.

With inhibition-dominated recurrent connectivity, our network is similar to an inhibition-stabilized network (ISN, 55). The hallmark of an ISN is a paradoxical effect: excitatory drive for inhibitory neurons may reduce their activity, and their inhibitory drive may increase it, due to recurrent effects. However, because we have not shown how to implement our model using inhibitory and excitatory neurons, it is unclear if our model will show such behavior. Interestingly, optogenetic perturbation experiments in the hippocampus report this paradoxical effect in CA1 and CA3 (56). The predictions of the ISN model also match experimental results from the cortex (57).

Our network model differs from the Stabilized Supralinear Network (SSN, 58, 59) when an exponent *n* = 1 is used in our model, but an exponent of *n* = 1.2 seems consistent with both models. This exponent can be measured experimentally by increasing an external tonic drive and measuring the response curve. For ISN, the response grows linearly, while for SSN it has both linear and sublinear regimes; for a tightly balanced E-I network, a phase transition occurs at some value of the external input in which the network becomes entrained by it and loses its asynchronous state.

### Limitations of the study

Our theoretical model does not capture biological heterogeneity because it assumes a single set of parameters across the population. The first kind of such heterogeneity is due to multiple neuronal populations, which may have distinct single-neuron properties. The second kind of heterogeneity is to allow for within-population variability in single-neuron properties. Our model-fitting results on pyramidal neurons in area CA3 suggest that a distribution of exponents may be a more realistic model, which we postpone for future work.

## ACKNOWLEDGEMENTS

This work was supported by the Wellcome Trust (Investigator Award in Science 212262/Z/18/Z to M.L.), the Human Frontiers Science Programme (Research Grant RGP0044/2018 to M.L.), and the Blavatnik Cambridge Postdoctoral Fellowships (to U.C.). We thank Peter Jonas, Christophe Mulle, Meryl Malézieux, Jeff Magee, Yiding Li, and May-Britt Moser for sharing their data. We thank Peter Jonas, Alessandro Treves, Yasser Roudi, Samuel Eckmann, Yiding Li, and Ashley Kees for useful discussions. We thank Christophe Mulle and Ashley Kees for sharing their spike detection and subthreshold voltage extraction code.

## DATA AND CODE AVAILABILITY

Requests for resources should be directed to Peter Jonas. Requests for additional information should be directed to the lead contact.

- All data reported in this paper will be shared upon request.
- All original code has been deposited at a public repository and will be publicly available as of the date of publication.
- Any additional information required to re-analyze the data reported in this paper is available from the lead contact upon request.

## Supplementary figures

**Fig. S1.**
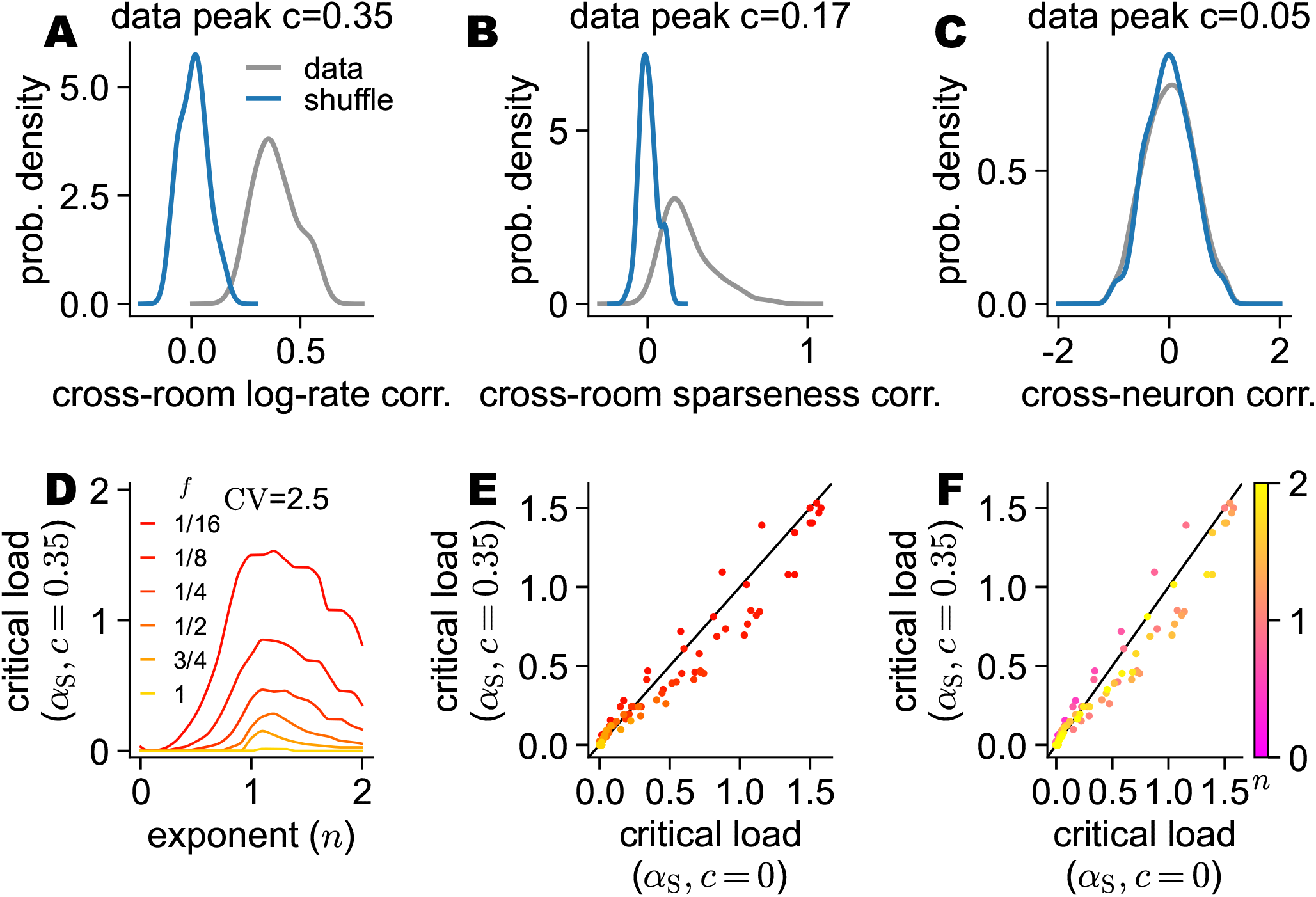
Theoretical results for correlated patterns. (A-C) Analysis of neural data from *Alme et al*., *2014* (15). (A) Distribution of correlation coefficients when comparing the log-rate between different rooms, using only neurons that are active in both rooms (all 55 possible comparisons). Data (gray) or shuffled neurons control (blue). (B) Distribution of correlation coefficients when comparing the binarized activity between different rooms (all 55 possible comparisons). Data (gray) or shuffled neurons control (blue). (C) Distribution of correlation coefficients when comparing the log-rate between different neurons, using only rooms in which both neurons are active (1000 random pairs). Data (gray) or shuffled neurons control (blue). (D) Simulation results for the critical load for stability, *α*_S_, for correlated patterns with *c* = 0.35 at different choices of the exponent *n* (x-axis) and sparseness *f* values (color-coded). Using a fixed pattern CV (value indicated) and the optimal threshold *θ*. (E-F) Scatter plots comparing the simulation results with *c* = 0.35 (D) and *c* = 0 Fig. 1F, colored by sparseness *f* (E) or exponent *n* (F).

**Fig. S2.**
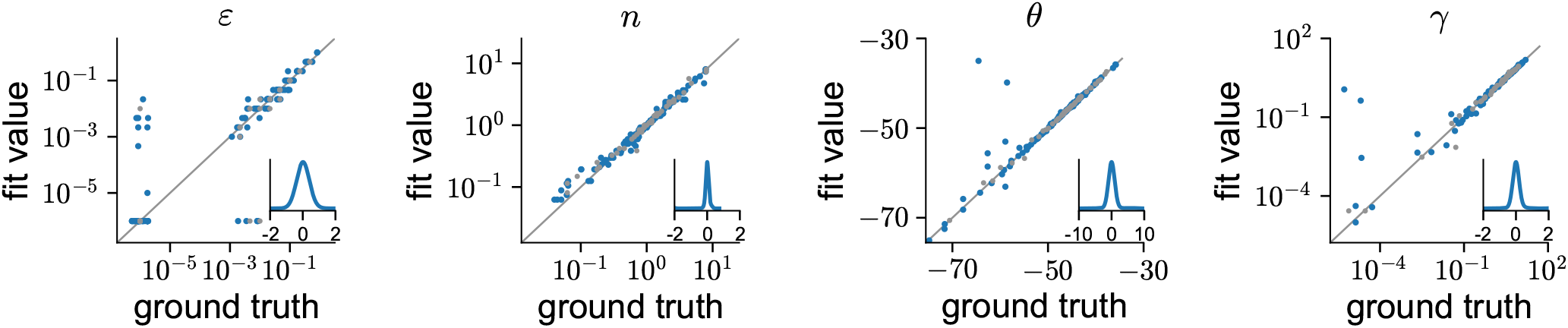
Parameter identification on synthetic data. Each panel presents a scatter plot comparing ground-truth values of a single parameter (indicated in the title) with the maximum-likelihood value found using the procedure described in ‘Fitting a Poisson process spiking model’. Each point is a result of an experiment where the real subthreshold voltage from a single neuron was used, but spike times were sampled from an inhomogeneous Poisson process model with an activation function defined by the four parameters from Eq. 2. The optimal values fitted to the neuron’s real spike times were used (gray dots) or a perturbed version of all four values (blue dots). Insets: histogram of the difference between the x and y coordinates (i.e., in linear scale for the threshold *θ* and log scale for other parameters).

**Fig. S3.**
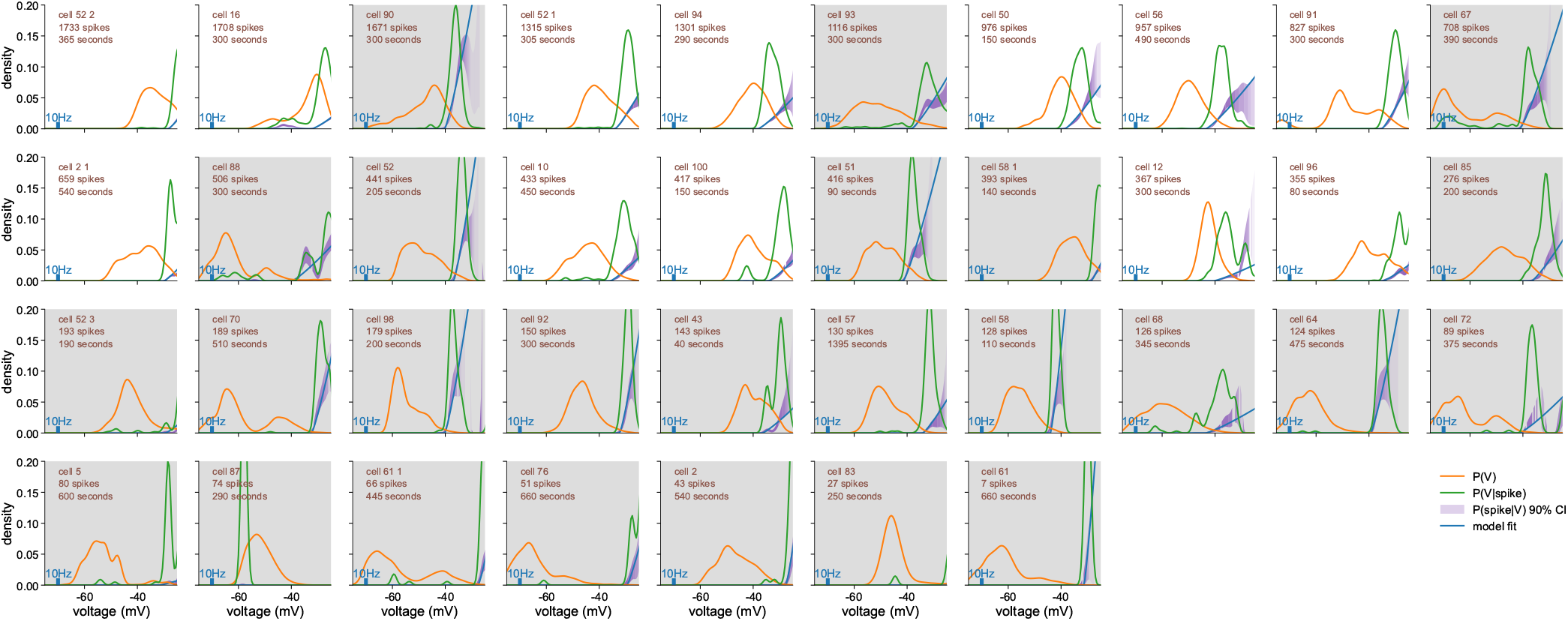
Analysis of voltage-dependent spiking probability in CA3 pyramidal neurons from *Malezieux et al*., *2020* (29). A panel for each intracellularly recorded cell, ordered by the number of spikes. Panel background color is white for ‘well-constrained’ cells, and gray for ‘poorly-constrained’ cells. Each panel presents the distribution of subthreshold voltage (orange), the distribution of voltage at spike initiation (green), and a 90 % credible interval for the resulting spike probability at each voltage (purple; lighter shade indicates a wider interval). The blue curve is the activation function of the best-fit parametric model, Eq. 2, in units of Hz (scale indicated at each panel) or in units of spike probability at each voltage (regular y-axis). The best-fit activation function has *n*\* = 1.19 and *ε*\* = 0.10 shared across all neurons; *θ*\* and *γ*\* were fit for each neuron.

**Fig. S4.**
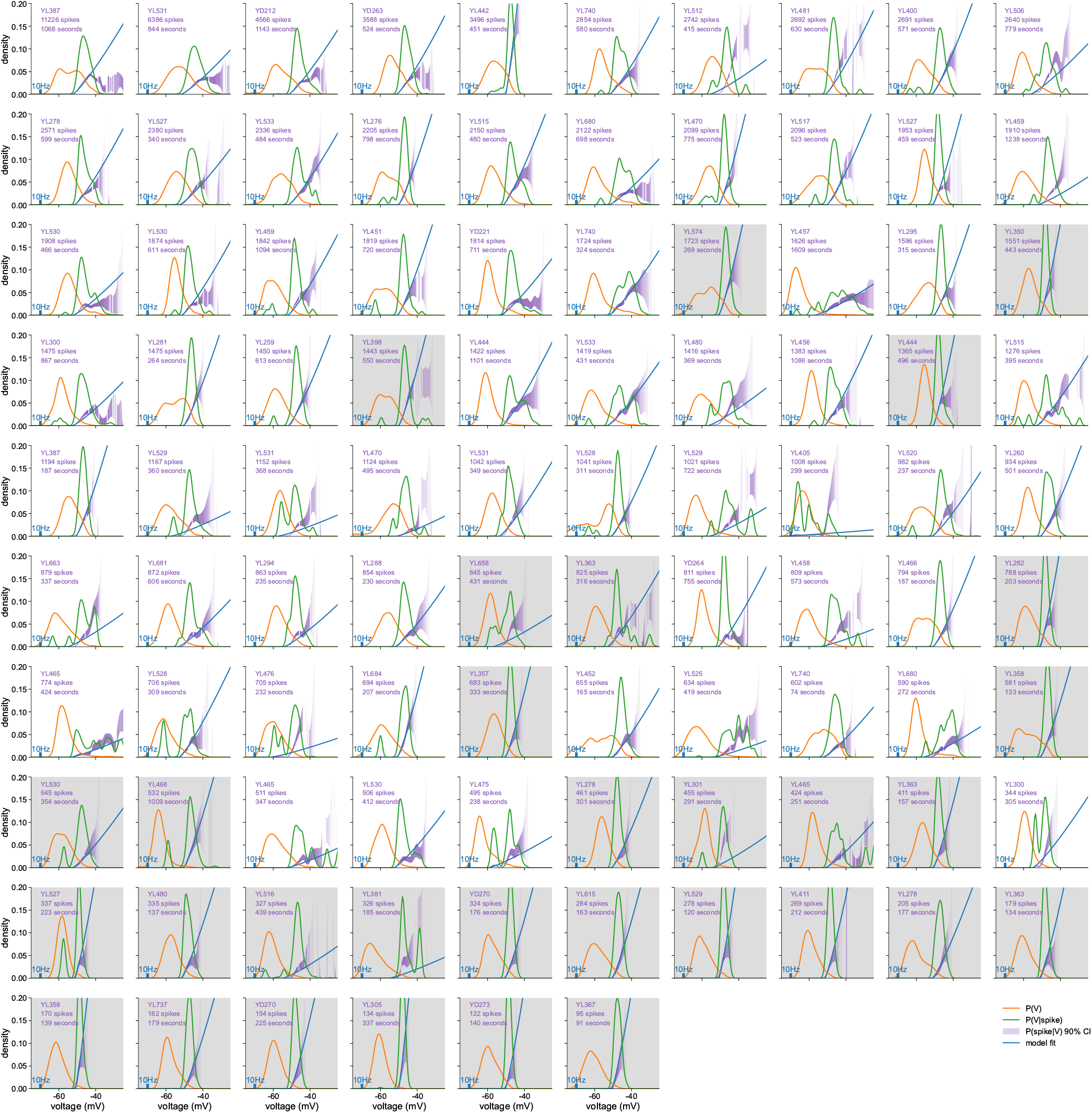
Analysis of voltage-dependent spiking probability in CA3 pyramidal neurons from *Li et al*., *2024* (8) A panel for each intracellularly recorded cell, ordered by the number of spikes. Panel background color is white for ‘well-constrained’ cells, and gray for ‘poorly-constrained’ cells. Each panel presents the distribution of subthreshold voltage (orange), the distribution of voltage at spike initiation (green), and a 90 % credible interval for the resulting spike probability at each voltage (purple; lighter shade indicates a wider interval). The blue curve is the activation function of the best-fit parametric model, Eq. 2, in units of Hz (scale indicated at each panel) or in units of spike probability at each voltage (regular y-axis). The best-fit activation function has *n*\* = 1.19 and *ε*\* = 0.10 shared across all neurons; *θ*\* and *γ*\* were fit for each neuron.

**Fig. S5.**
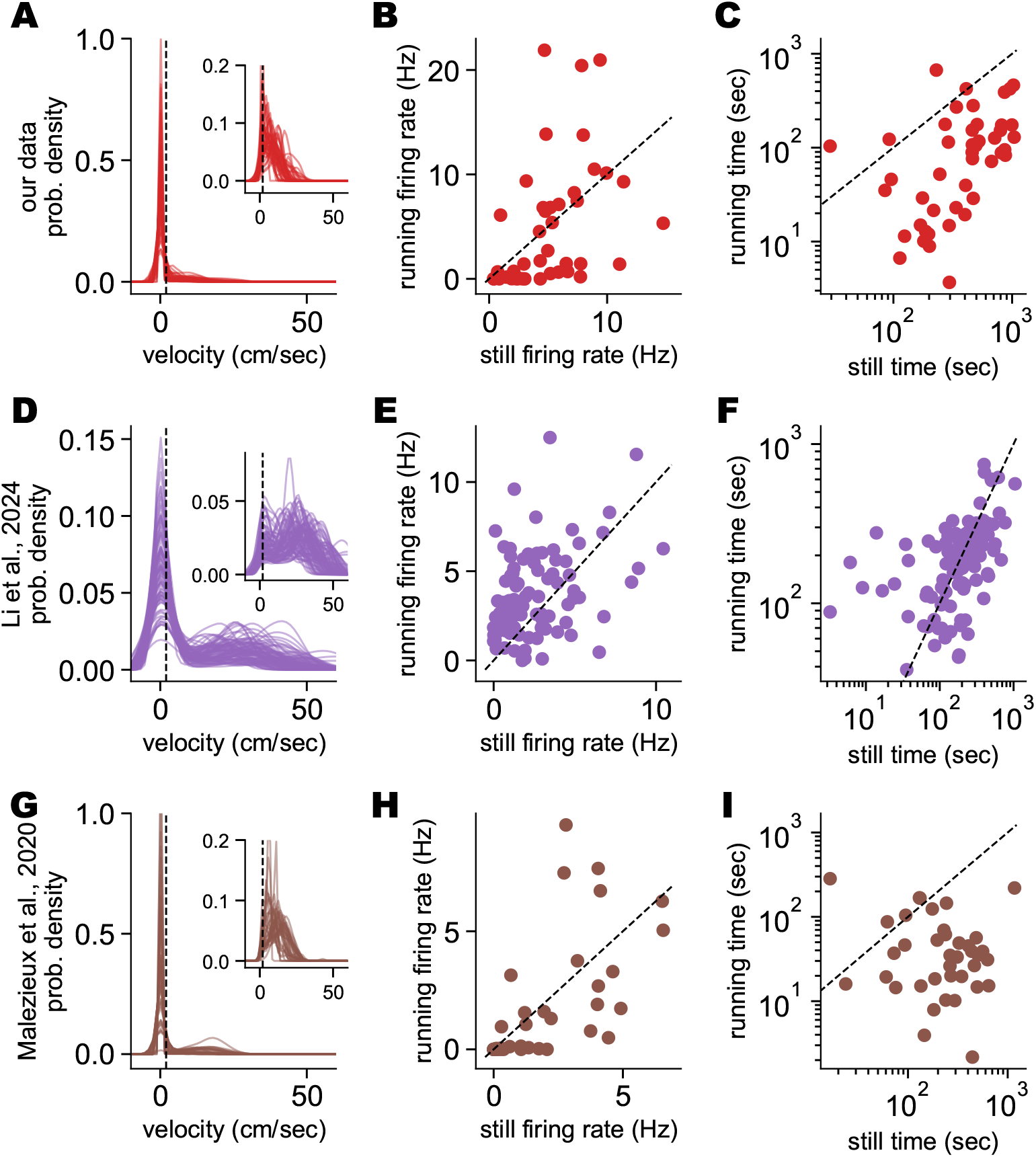
Dataset differences with respect to velocity and behavioral state. Each rows depict the statistics of a different dataset, as indicated in the leftmost panel. (A,D,G) Velocity density of each animal. Inset: the same for velocities with absolute values above 0.5 cm/sec. Dashed vertical line indicates the running threshold used in behavioral analysis. (B,E,H) Scatter plot comparing firing rate when the animal is in the ‘still’ state (x-axis) versus the ‘running’ state, for each animal. (C,F,I) Scatter plot comparing the time spent by the animal in the ‘still’ state (x-axis) versus the ‘running’ state, for each animal.

**Fig. S6.**
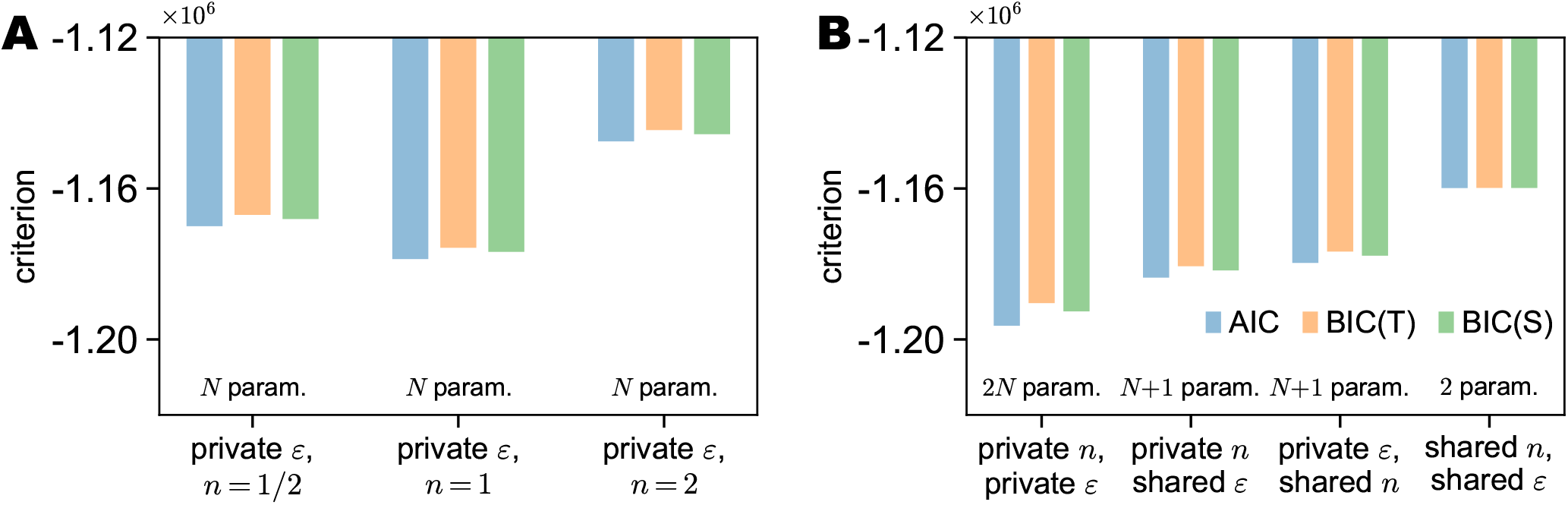
Model selection. Three model-comparison criteria (AIC, BIC(T), and BIC(S); see Methods) applied to different models (x-axis). These methods combine the models’ log-likelihood with the number of fitted parameters; lower is better. The number of parameters of each model is indicated at the bottom. (A) Comparing “point models” with private noise *ε* and a point value for the exponent *n*. (B) Comparing the four models defined by a private or shared noise *ε* and a private or share exponent *n*.

## Methods

### Simulation results of the critical load for stability

Memory patterns were assumed to be sparse, with a fraction 0 *< f* ≤ 1 of the neural population active in each pattern, and (1 − *f*)*N* of the neurons are exactly zero. The non-zero activations are sampled i.i.d. (independently and identically distributed) from a log-normal distribution. Sampling from a log-normal distribution with meanJ*m*_0_ and standard deviation *s*_0_ is done taking *exp*(*G*) for samples from a Gaussian *G* ~ N (*m, s*^2^) where 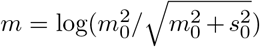 and 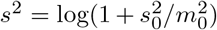. Sampling from a correlated log-normal distribution (where neuronal activity is correlated across patterns) with correlation *c* is done by sampling *G* as a sum of *P* uncorrelated patterns scaled by 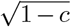 and a single shared pattern scaled by 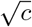. In our simulations, the log-normal distribution was parametrized by its coefficient of variation CV = *s*_0_*/m*_0_, while fixing its mean to be *m*_0_ = 1.

Memory patterns are stored in a network of *N* neurons with voltage dynamics: 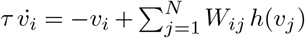 where *v*_*i*_ is the voltage of neuron *i, τ* is the neural time constant (assumed to be shared across neurons), **W** ∈ ℝ^*N* ×*N*^ defines synaptic connection weights (such that *W*_*ij*_ is the strength of the synaptic connection from neuron *j* to neuron *i*), and *h*(*v*), defined in Eq. 1, is the activation function that maps the ‘voltage’ of a neuron, *v*_*i*_, to its (positive) instantaneous firing rate (also assumed to be homogeneous across neurons).

The above voltage dynamics have equivalent rate dynamics (17), defined as 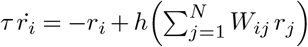. A pattern **r**_0_ would be a fixed point of the dynamics when **r**_0_ = **h** (**Wr**_0_), and a stable fixed point if the eigenvalues of the Jacobian of the rate dynamics at the pattern **J** = −**I** + diag *h*^′^ *h*^−1^(**r**_0_) **W** have only negative real parts.

For a set of *P* memory patterns **r**^*µ*^ for *µ* = 1 … *P*, we solved the problem of finding minimal-norm weights such that each memory pattern is a fixed-point of the network dynamics, without self-connections:

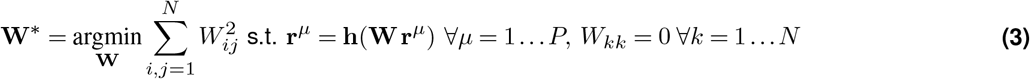

This is a convex problem with linear equality and inequality constraints (the inequalities result from the rectification of the activation function) and can be solved using off-the-shelf optimizers (60).

Assuming both the number of neurons *N* and the number of memory patterns *P* are large, the number of patterns that can be stored as *fixed points* of the network is *P*_max_ = *Nα*_C_, while the number of patterns that can be stored as *stable fixed points* is *P*_max_ = *Nα*_S_ (14). When the number of activity patterns is below the critical load for stability, the network dynamics would perform stable memory recall.

The value of *α*_S_ (*f*, CV, *θ, n*) is calculated in simulations using a large set of parameter combinations. These combinations were generated using a fixed grid *f* values (1*/*16, 1*/*8, 1*/*4, 1*/*2, 3*/*4, 1), CV values (1*/*8, 1*/*4, 1*/*2, 1, 2, 4, 8), and exponents *n* ∈ {0.1 *i* : *i* = 1 … 20}, and an adaptive grid of *θ* values. For each parameter combination, optimal weights **W** were obtained using Eq. 3 for a network of size *N* = 256. We started from *P* = 4, *θ* ∈ [− 6, 3] (with equal steps in linear scale), and at each iteration increased *P* by 1 and made the range of *θ* narrower, removing leading or trailing values without stable patterns, and re-adjusting grid steps. The numerical value of *α*_S_ reported for a given parameter combination was defined as *P/N* for the maximal value of *P* for which at least half of the patterns were dynamically stable.

We claim the two extra parameters in Eq. 2, a gain *γ* and a noise term *ε*, do not change the results compared with the model defined by Eq. 1. First, introducing a global gain *γ* and scaling the threshold as *θ*_*γ*_ = *γ*^−1*/n*^*θ* would yield, when solving Eq. 3, a scaled solution *W*_*γ*_ = *γ*^−1*/n*^*W*. Then, both the fixed-points equation and the Jacobian matrix are unmodified. Second, as long as *ε* is small, assuming the same pattern statistics but with *ε* added to each pattern, introduces only a small perturbation to the mean of the patterns and does not affect other moments, and thus would not affect the results. Second, as long as *ε* is small, assuming the pattern statistics are unmodified except for *ε* added to each activation introduces only a small perturbation to the mean of the patterns and does not affect other moments, and thus would not affect the results. Regarding heterogeneity, as with the threshold and exponent, our theoretical analysis does not currently handle per-neuron parameters (see Discussion).

### CA3 ongoing activity statistics

The neural data from *Alme et al*., *2014* (15) included the response of *N* = 332 neurons in 11 rooms, from which we calculated the firing rate of each neuron in each room. 5-25 % of the neurons were silent in each room. A probability density was created for the log_10_ firing rate of active neurons in each room by using KDE (scipy.stats.gaussian_kde in Python, Silverman method to determine width). A global optimal fit was found by calculating the mean and standard deviation of the log_10_ firing rate, pooling data across all rooms. The coefficient of variation was estimated directly from their (linear) firing rates. A cross-room firing rate correlation coefficient was calculated from the log_10_ firing rate for each pair of rooms, using all neurons active in both rooms. A cross-neuron firing rate correlation coefficient was calculated from the log_10_ firing rate for 1000 pairs of neurons, in all rooms where both were active. A cross-room sparseness correlation coefficient was calculated from binarized firing rate (i.e., 0 if the neuron never fires in this room and 1 otherwise) for each pair of rooms.

### Animal experiments

Whole-cell patch-clamp recordings *in vivo* were performed in 63-to 224-day-old C57BL/6J mice (RRID:IMSR_JAX:000664). Animals were housed under a reversed light cycle (dark: 7:00 am – 7:00 pm, light: 7:00 pm – 7:00 am). For experiments, both male and female animals were used. All experiments were carried out in strict accordance with institutional, national, and European guidelines for animal experimentation and approved by the Austrian Bundesministerium für Wissenschaft, Forschung und Wirtschaft (BMWFW-66.018/0007-WF/II/3b/2014) and Bundesmin-isterium für Bildung, Wissenschaft und Forschung (BMWFW-66.018/0010-V/3b/2019).

### Surgery and animal training

Head-bar implantation and craniotomies were performed under anesthesia induced via in-traperitoneal (i.p.) injection of 80 mg/kg ketamine (Intervet GesmbH, Vienna, Austria) and 8 mg/kg xylazine (Dr. E. Graeub AG, Bern, Switzerland), supplemented with constant oxygen supply. Following local anesthesia with lidocaine (AstraZeneca Österreich GmbH, Vienna, Austria), the skull was exposed and cleaned. A small craniotomy (500 µm diameter) was drilled above the cerebellum to position a gold pin, which was secured with tissue glue (Vetbond, 3M Austria GmbH, Vienna, Austria) to serve as a grounding electrode for multichannel recordings.

The skull was subsequently covered with a thin layer of UV-curing All-In-One OptiBond adhesive agent (Kerr Corporation, Orange, CA, USA). A sterilized head post was fixed to the coated skull using UV-curing optical adhesive (Charisma, Kulzer GmbH, Hanau, Germany), followed by an anchoring layer of dental cement (Paladur; Heraeus Holding GmbH, Hanau, Germany). During the application of the final dental cement layer, the surface was smoothed to prevent post-operative scalp irritation. Post-operative analgesia was maintained with i.p. injections of 4 mg/kg meloxicam (Boehringer Ingelheim RCV GmbH, Vienna, Austria) and 0.15 mg/kg buprenorphine twice daily for two days post-surgery. Mice were single-housed and provided with a nutrient-rich diet during a one-week recovery period.

Five days post-surgery, home cages were enriched with running wheels and tunnels. One week post-surgery, mice were placed on a restricted water schedule (1.2 ml/day on weekdays; ad libitum water access over weekends). After one week of habituation to this water restriction regime, the daily volume was reduced to 1 ml/day, and mice were introduced to a linear belt training apparatus. On the first day, animals freely explored the apparatus for 5 min. On the second day, mice were head-fixed but not tightly restrained within the setup for 10 min. Regular training sessions commenced on the third day, during which mice were trained to run a distance of 180 cm to receive a water reward for 30 min. Training continued for 7–10 days until mice demonstrated frequent and stable running behavior across three consecutive days.

One day prior to the recording experiments, two small craniotomies (0.5 mm diameter) were gently drilled bilaterally for intracellular and extracellular recordings. Stereotaxic coordinates for intracellular recordings were 1.7 mm mediallateral (ML, from the midline) and 1.5 mm anterior-posterior (AP, from bregma). Coordinates for local field potential (LFP) recordings were 1.7–1.9 mm ML and 2.6 mm AP.

### Behavioral set-up

A custom-made linear belt system was utilized for electrophysiological recordings as described previously (44). Animals ran on a 180 cm black velvet fabric belt (McMaster-Carr, Elmhurst, IL, USA), divided into equal lengths enriched with three distinct tactile textures made of white hot-glue and white Velcro: Velcro strips, glued dots, and curved zig-zag tracks. Two large Styrofoam objects were added to each texture zone.

Water rewards were delivered upon entry into the Velcro strip zone via a transistor-transistor logic (TTL) pulse sent to a syringe pump. Anticipatory and consummatory licking behavior was detected using an optical sensor that measured the disruption of an infrared (IR) light beam (RS Components Handelsges.m.b.H., Gmünd, Austria). Spatial transitions to new textures and the completion of full belt rounds were detected by IR reflective sensors (RS Components Handelsges.m.b.H., Gmünd, Austria). The animal’s running speed and spatial location were monitored via an incremental rotary encoder positioned on one axis of the running wheel. An Arduino (Arduino S.r.l., Monza, Italy)-based interface converted these inputs into analog signals representing velocity, location, current texture, and lick detection. All analog behavioral signals were routed to the analog-to-digital (A/D) inputs of a HEKA amplifier (HEKA Elektronik GmbH, Lambrecht, Germany) to ensure precise temporal synchronization with the electrophysiological recordings.

### In vivo electrophysiology

LFPs were recorded using a single-shank silicon probe designed for acute recordings, featuring 32 recording channels with electrodes 50 µm apart spanning the entire depth of the hippocampus (15 µm thickness, A1×32; NeuroNexus Inc., Ann Arbor, MI, USA). The probe was connected via an OMNETICS adapter to an Intan stimulation and recording system controlled by customized Intan software (Intan Technologies, Los Angeles, CA, USA). Following a micro-incision into the dura, the silicon probe was advanced through the neocortex at a speed of 10 µm/s. Before reaching the hippocampus, the advancement speed was reduced to 1 µm/s until the probe entered the CA3 region (dorso-ventral depth: 2200 µm).

Whole-cell patch-clamp measurements in awake mice were performed according to previously established protocols (40, 44). Patch pipettes were fabricated using a horizontal Brown-Flaming micropipette puller (P-1000; Sutter Instrument Company, Novato, CA, USA) from borosilicate glass capillaries (Karl Hilgenberg GmbH, Malsfeld, Germany; 1.75 mm outer diameter, 1.25 mm inner diameter). The resulting pipettes exhibited a long, thin taper (7 µm in diameter at a distance of 100 µm from the tip) with a tip resistance of 5–8 MΩ. Pipettes were filled with an intracellular solution containing 130 mM K-gluconate, 2 mM KCl, 10 mM HEPES, 2 mM MgCl2, 2 mM Na2ATP, 0.3 mM NaGTP, 18 mM sucrose, 0.1 mM EGTA, and 0.3% biocytin for post-hoc morphological identification (pH adjusted to 7.3 with KOH). To prevent pipette clogging, a positive pressure of 700 mbar was maintained until the dura was penetrated and was immediately reduced to 400 mbar as the pipette advanced through the neocortex. After traversing the neocortex, the positive pressure was lowered to 180 mbar, and finally to 20–25 mbar at a depth of 1800 µm. The approach toward a putative cell body was detected by a consistent increase in pipette resistance. Mild suction (−25 mbar) was then applied to form a high-resistance seal onto the cell. Following a 1–2-minute equilibration period to clear excessive extracellular K+ ions, a brief, strong suction pulse was applied to establish whole-cell configuration. A test voltage pulse (stepping to +50 mV and −10 mV) was recorded to identify the neuron and determine the series resistance. Recordings with a series resistance exceeding 100 MΩ and less than 5-minute recording time were discarded. Following pipette capacitance correction (100%) and bridge-balance compensation in the whole-cell configuration, all signals were sampled at 25 kHz and low-pass filtered at 10 kHz. Active and passive membrane properties of CA3 pyramidal neurons were evaluated using a current-injection protocol ranging from −100 to 400 pA in 50 pA increments. Spontaneous activity was recorded with zero current injection throughout the baseline session. All intracellular data were acquired using an EPC 10 double amplifier (HEKA Elektronik GmbH, Lambrecht, Germany) controlled by PatchMaster acquisition software (v2×90.1). Data synchronization of the two recording systems (Intan and Heka) was ensured by a common trigger signal, recorded by both systems, and offline synchronization of both data sets. After completing the recordings, the patch pipettes were slowly retracted to allow outside-out patch formation for subsequent post-hoc CA3 cell labeling.

### Subthreshold voltage extraction

Electrophysiology results from *N* = 49 putative CA3 pyramidal neurons were processed to analyze the relation between the neuron’s voltage and spiking activity. Voltage traces from the neurons were manually observed to identify whether voltage levels at the beginning or the end of the experiment are distinct from those in the middle of the experiment. Following this analysis, voltage traces from the beginning of the experiment were removed for five of the neurons, and at the end of the experiment for seven of the neurons. The voltage trace was down-sampled from 25000 Hz to 12500 Hz, prior to spike time identification and extraction of subthreshold voltage.

#### Spike initiation

Spike initiation times were found by calculating the voltage derivative ‘dVdt’ (V/sec) through finite differentiation, and candidate times were all time points where ‘dVdt’ intersected ‘dVdt threshold’ (5 V/sec) from below. These were filtered to remove times that occur within ‘refractory period’ (0.5 ms) of previous ones, remove times that are not followed by a voltage peak within a ‘peak window’ (5 ms), and remove times that are followed by the same peak as previous ones. Peaks were defined through a signal processing algorithm (scipy.signal.find_peaks in Python, with ‘prominence’ of 1 and ‘distance’ defined by the refractory period), and were filtered to leave peaks above ‘Vm threshold’ (−25 mV).

#### Spike termination

The spike termination time is found by calculating the zero crossings of ‘dVdt’ where its slope (i.e., the second voltage derivative) is positive. For each spike, if such zero crossings were found within a specific time window, this value was the spike termination; otherwise, the time point with the minimal absolute ‘dVdt’ within this window is considered the spike termination. This time window was defined by starting from a spike’s (last) peak and finding the minimal ‘dVdt’ (i.e., largest decrease) within one refractory period (0.5 ms) from that peak. Then the time window was between the time point with the minimal ‘dVdt’ and the early of the initiation time of the following spike, and 5 ms after the peak.

#### Subthreshold voltage

Spike times were the times of spike initiation (at precision defined by the 12500 Hz sampling), and an initial version of the subthreshold voltage was created by interpolating the voltage at time points between spike initiation and termination. The full and subthreshold voltages were then subsampled to 1250 Hz (without changing spike-time precision). The final subthreshold voltage is created from the initial one by finding, for each spike (except those affected by this analysis for previous spikes), the first place within the search window (2 sec) where voltage falls below the spike threshold, and removing intermediate voltages. If voltage stays above the threshold within the search window, intermediate voltages are discarded for the entire window. This means that the voltage of all the spikes in a burst is assumed to be the same as the voltage of its first spike.

The same procedure was applied to our data and data from *Malezieux et al*., *2020* (29), *Li et al*., *2024* (8).

### Analysis of animal state

Behavioral information, such as position and velocity, was collected at a rate of 2500 Hz. For the identification of animal state (‘running’ versus ‘still’), velocity was smoothed using a moving average with windows length of 0.5 sec. Running periods were defined as times when velocity exceeded a threshold of 2 cm/sec, merging subsequent periods that are less than 3 sec apart (from offset to onset), and finally discarding periods of less than 0.5 sec. All periods not classified as ‘running’ were considered as ‘still’.

Both the non-parametric analysis of voltage-dependent spike probability and the parametric analysis of fitting a Poisson process model, which were performed on the entire voltage trace, were repeated on voltage and spike times from either all ‘running’ periods or all ‘still’ periods.

The same procedure was applied to our data and data from *Malezieux et al*., *2020* (29), *Li et al*., *2024* (8), with the caveat that the behavioral information collected there differed in sampling rate.

### Voltage-dependent spike probability

At a sampling rate of 1250 Hz, we assume spiking is a binary event that occurs or does not occur at any given time. Time points where voltage was below −100*mV* or above 0*mV* were discarded from the analysis, along with spikes occurring at those times. A non-parametric Bayesian approach was used to calculate the calculation of spiking probability dependent on voltage, through application of Bayes’s rule:

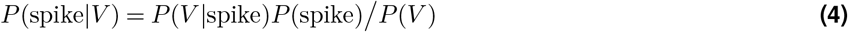

If voltage is assumed to be digitized into non-overlapping bins, we can directly count *C*(*V*|spike), the number of voltage visits to *V* during a spike, and *C*(*V*), the total number of visits to *V*. Then, denoting *T* as the total number of time points and *S* the total number of spikes, the above probabilities can be calculated as *P* (*V*|spike) = *C*(*V*|spike)*/S* and *P* (*V*) = *C*(*V*)*/T*. This approach is straightforward, but may create artifacts from the arbitrary choice of bin edges. To avoid this limitation, we estimate *P* (*V*) from the voltage at all time points using Kernel Density Estimation (KDE) with a width of 1*mV* (using the FFTKDE algorithm of the KDEpy Python package). Similarly, *P* (*V*|spike) is estimated from the voltage at time points where a spike occurred, and *P* (spike) is the fraction of time points where spiking occurred. The temporal binning used was *dt* = 1*/*1250*sec*.

#### Credible interval

The resulting estimate *P* (spike|*V*) is a function of the voltage, but otherwise we have not assumed it takes any specific shape; for example, it is not guaranteed to be monotonically increasing. Furthermore, this is a noisy statistic, with possible sources of stochasticity coming from both the neuron (e.g., having a noisy spike threshold) and the experimental protocol (e.g., noisy measurements of voltage), and the amount of noise in this estimate depends on the finite number of measurements. Thus, we are not interested in the point estimate of *P* (spike|*V*) recovered by Bayes’s rule, but rather in a credible interval for it. We suggest that a credible interval for this quantity can be estimated by assuming that *P* (spike|*V*) ~ *Beta*(*α, β*), namely Beta-distributed with *α* = *C*(spike|*V*) and *β* = *C*(no spike|*V*) so that *α* + *β* = *C*(*V*). We thus report a 90% credible interval for *P* (spike *V*) as the interval given by the 5% and 95% percentiles of this distribution, which is an analytic value (scipy.stats.beta.ppf in Python). This estimate depends on the binning of time *dt* only through the assumption that binning is fine enough that spiking becomes a binary event at each time point, and does not require binning of the voltage *dV*, beyond the choice of the width parameter for KDE, as it can be calculated for any *V* without binning given the KDE estimators. Those results were deemed ‘well-constrained’ if the 90 % credible interval lower bound was above 0.001 and the width (difference between upper and lower bound) was less than 0.01 for at least 5 mV, and ‘poorly-constrained’ otherwise.

#### Historical note

Our method is similar to methods developed to analyze the relation between voltage and firing rate in neuronal responses of visual areas (32, 33). These methods extract subthreshold voltage and combine it with spike times to estimate the firing-rate distribution given voltage *P* (*r*|*V*). The classic method low-pass-filtered subthreshold voltage to 24 Hz, and spikes were converted into firing rate with the same temporal resolution; then the conditional probability *P* (*r*|*V*) was estimated through discrete binning of the voltage (32). The more recent version binned both voltage (into 1*mV* bins) and time (into 20*ms* bins), then estimated the conditional probability *P* (*r*|*V*) as we do. Our method replaced voltage binning with a KDE kernel of the same width and required neither temporal binning nor low-pass filtering. Operating at 1250 Hz, each time point either has a spike or not, which simplifies notation.

The same procedure was applied to our data and data from *Malezieux et al*., *2020* (29), *Li et al*., *2024* (8).

### Fitting a Poisson process spiking model

For a Poisson process generating spikes with voltage-dependent rate *r*(*V*), so that the spike probability dependence on the voltage is given by *P* (spike|*V*) = *dtr*(*V*), the log-likelihood of an entire voltage trajectory 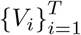 with *S* spikes triggered at voltages 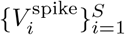 is given by:

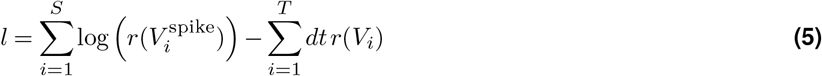

By assuming a specific parametric model for *r*(*V*), its maximum-likelihood parameters can be found through direct optimization. We used the parametric model from Eq. 2, where *ε* is a small number representing the baseline of voltage-independent spiking (required to prevent infinite values in the log-likelihood), *γ* is a gain factor, *θ* is a threshold, and *n* is an exponent. In our analysis, we usually assume that *ε* and *n* are shared across neurons, while *θ* and *γ* need to be optimized independently for each neuron.

The same procedure was applied to our data and data from *Malezieux et al*., *2020* (29), *Li et al*., *2024* (8).

### Statistical model comparison

Three criteria were used for model comparison. AIC (61) is defined as AIC = 2*k* − 2 log *L* where *k* is the number of model parameters and *L* is the model’s likelihood. BIC (62) is defined as BIC = *k* log *M* − 2 log *L* where *M* is the number of data points. We used two variants, BIC(*S*) where *M* = |*S*| is the number of spikes or BIC(*T*) where *M* = |*V*| is the number of voltage data points.

While Eq. 2 defines a model with four parameters, we suggest that two of them, the threshold *θ* and gain *γ*, differ substantially between neurons and thus need to be fit separately for each neuron. For a dataset of *N* neurons, we consider four classes of models which can be used to explain the data: (i) private exponent *n*, and noise *ε*, or *k* = 2*N* degrees of freedom; (ii) private exponent *n*, and shared noise *ε*, or *k* = *N* + 1 degrees of freedom; (iii) shared exponent *n*, and private noise *ε*, or *k* = *N* + 1 degrees of freedom; (iv) shared exponent *n*, and noise *ε*, or *k* = 2 degrees of freedom. We also consider “point models” with shared noise *ε* where the exponent is kept fixed at 1*/*2, 1, or 2, which have *k* = *N* degrees of freedom. See Fig. S6.

### Synthetic data control

For each neuron in our dataset, four “synthetic traces” were created by using the real subthreshold voltage and synthetic spike times, drawn from a Poisson point process with inhomogeneous rate 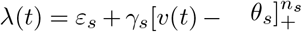, where *v*(*t*) is the neuron’s subthreshold voltage. In one trace for each neuron, the four parameters used were the optimal parameters found: *ε*_*s*_ = *ε*\*, *n*_*s*_ = *n*\*, *γ*_*s*_ = *γ*\*, *θ*_*s*_ = *θ*\*. In three traces for each neuron, the four parameters used were noisy versions of the optimal values, created as 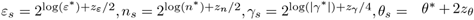, where *z*_*ε*_, *z*_*n*_, *z*_*γ*_, *z*_*θ*_ are i.i.d standard Gaussian variables. Those traces were then analyzed using the above ‘Fitting a Poisson process spiking model’, in order to check how well we recover the ground truth parameters.

## Bibliography

1. David Marr. Simple memory: a theory for archicortex. Philosophical Transactions of the Royal Society of London. B, Biological Sciences, 262(841):23–81, 1971.

2. Bruce L McNaughton and Richard GM Morris. Hippocampal synaptic enhancement and information storage within a distributed memory system. Trends in neurosciences, 10(10): 408–415, 1987.

3. Alessandro Treves and Edmund T Rolls. Computational analysis of the role of the hippocampus in memory. Hippocampus, 4(3):374–391, 1994.

4. Kazu Nakazawa, Michael C Quirk, Raymond A Chitwood, Masahiko Watanabe, Mark F Yeckel, Linus D Sun, Akira Kato, Candice A Carr, Daniel Johnston, Matthew A Wilson, et al. Requirement for hippocampal ca3 nmda receptors in associative memory recall. science, 297(5579):211–218, 2002.

5. Tom J Wills, Colin Lever, Francesca Cacucci, Neil Burgess, and John O’Keefe. Attractor dynamics in the hippocampal representation of the local environment. Science, 308(5723): 873–876, 2005.

6. Takuya Sasaki, Norio Matsuki, and Yuji Ikegaya. Metastability of active ca3 networks. Journal of Neuroscience, 27(3):517–528, 2007.

7. Joshua P Neunuebel and James J Knierim. Ca3 retrieves coherent representations from degraded input: direct evidence for ca3 pattern completion and dentate gyrus pattern separation. Neuron, 81(2):416–427, 2014.

8. Yiding Li, John J Briguglio, Sandro Romani, and Jeffrey C Magee. Mechanisms of memory-supporting neuronal dynamics in hippocampal area ca3. Cell, 187(24):6804–6819, 2024.

9. John J Hopfield. Neural networks and physical systems with emergent collective computational abilities. Proceedings of the national academy of sciences, 79(8):2554–2558, 1982.

10. MV Tsodyks. Associative memory in asymmetric diluted network with low level of activity. Europhysics Letters, 7(3):203, 1988.

11. MR Bennett, WG Gibson, and J Robinson. Dynamics of the ca3 pyramidial neuron autoassociative memory network in the hippocampus. Philosophical Transactions of the Royal Society of London. Series B: Biological Sciences, 343(1304):167–187, 1994.

12. Segundo Jose Guzman, Alois Schlögl, Michael Frotscher, and Peter Jonas. Synaptic mechanisms of pattern completion in the hippocampal ca3 network. Science, 353(6304):1117–1123, 2016.

13. Jake F Watson, Victor Vargas-Barroso, and Peter Jonas. Cell-specific wiring routes information flow through hippocampal ca3. Cell Reports, 44(8), 2025.

14. Uri Cohen and Máté Lengyel. Dynamical stability for dense patterns in discrete attractor neural networks. arXiv preprint arXiv:2507.10383, 2025.

15. Charlotte B Alme, Chenglin Miao, Karel Jezek, Alessandro Treves, Edvard I Moser, and May-Britt Moser. Place cells in the hippocampus: eleven maps for eleven rooms. Proceedings of the National Academy of Sciences, 111(52):18428–18435, 2014.

16. György Buzsáki and Kenji Mizuseki. The log-dynamic brain: how skewed distributions affect network operations. Nature Reviews Neuroscience, 15(4):264–278, 2014.

17. Kenneth D Miller and Francesco Fumarola. Mathematical equivalence of two common forms of firing rate models of neural networks. Neural computation, 24(1):25–31, 2012.

18. John J Hopfield. Neurons with graded response have collective computational properties like those of two-state neurons. Proceedings of the national academy of sciences, 81(10): 3088–3092, 1984.

19. Kenji Mizuseki and György Buzsáki. Preconfigured, skewed distribution of firing rates in the hippocampus and entorhinal cortex. Cell reports, 4(5):1010–1021, 2013.

20. I Kanter and Haim Sompolinsky. Associative recall of memory without errors. Physical Review A, 35(1):380, 1987.

21. Elizabeth Gardner. The space of interactions in neural network models. Journal of physics A: Mathematical and general, 21(1):257, 1988.

22. Omar J Ahmed and Mayank R Mehta. The hippocampal rate code: anatomy, physiology and theory. Trends in neurosciences, 32(6):329–338, 2009.

23. Jake F Watson, Victor Vargas-Barroso, Rebecca J Morse-Mora, Andrea Navas-Olive, Mojtaba R Tavakoli, Johann G Danzl, Matthias Tomschik, Karl Rössler, and Peter Jonas. Human hippocampal ca3 uses specific functional connectivity rules for efficient associative memory. Cell, 188(2):501–514, 2025.

24. Frances S Chance, Larry F Abbott, and Alex D Reyes. Gain modulation from background synaptic input. Neuron, 35(4):773–782, 2002.

25. Zachary F Mainen and Terrence J Sejnowski. Reliability of spike timing in neocortical neurons. Science, 268(5216):1503–1506, 1995.

26. Michael N Shadlen and William T Newsome. Noise, neural codes and cortical organization. Current opinion in neurobiology, 4(4):569–579, 1994.

27. Carl Van Vreeswijk and Haim Sompolinsky. Chaos in neuronal networks with balanced excitatory and inhibitory activity. Science, 274(5293):1724–1726, 1996.

28. A Aldo Faisal, Luc PJ Selen, and Daniel M Wolpert. Noise in the nervous system. Nature reviews neuroscience, 9(4):292–303, 2008.

29. Meryl Malezieux, Ashley L Kees, and Christophe Mulle. Theta oscillations coincide with sustained hyperpolarization in ca3 pyramidal cells, underlying decreased firing. Cell reports, 32(1), 2020.

30. Yoav Adam, Jeong J Kim, Shan Lou, Yongxin Zhao, Michael E Xie, Daan Brinks, Hao Wu, Mohammed A Mostajo-Radji, Simon Kheifets, Vicente Parot, et al. Voltage imaging and optogenetics reveal behaviour-dependent changes in hippocampal dynamics. Nature, 569 (7756):413–417, 2019.

31. Qixin Yang, Shulamit Baror-Sebban, Rotem Kipper, Michael London, and Yoav Adam. Alloptical electrophysiology reveals behavior-dependent dynamics of excitation and inhibition in the hippocampus. Neuron, 114(9):1635–1650, 2026.

32. Matteo Carandini and David Ferster. Membrane potential and firing rate in cat primary visual cortex. The Journal of Neuroscience, 20(1):470–484, 2000.

33. Merse E Gáspár, Pierre-Olivier Polack, Peyman Golshani, Máté Lengyel and Gergő Orbán. Representational untangling by the firing rate nonlinearity in v1 simple cells. Elife, 8:e43625, 2019.

34. Ivan Cohen and Richard Miles. Contributions of intrinsic and synaptic activities to the generation of neuronal discharges in in vitro hippocampus. The Journal of physiology, 524(Pt 2):485, 2000.

35. Janina Kowalski, Jian Gan, Peter Jonas, and Alejandro J Pernía-Andrade. Intrinsic membrane properties determine hippocampal differential firing pattern in vivo in anesthetized rats. Hippocampus, 26(5):668–682, 2016.

36. Qian Sun, Alaba Sotayo, Alejandro S Cazzulino, Anna M Snyder, Christine A Denny, and Steven A Siegelbaum. Proximodistal heterogeneity of hippocampal ca3 pyramidal neuron intrinsic properties, connectivity, and reactivation during memory recall. Neuron, 95(3):656–672, 2017.

37. DA Turner, X-G Li, GK Pyapali, A Ylinen, and G Buzsaki. Morphometric and electrical properties of reconstructed hippocampal ca3 neurons recorded in vivo. Journal of Comparative Neurology, 356(4):580–594, 1995.

38. W Alden Spencer and Eric R Kandel. Hippocampal neuron responses to selective activation of recurrent collaterals of hippocampofugal axons. Experimental Neurology, 4(2):149–161, 1961.

39. Christopher D Harvey, Forrest Collman, Daniel A Dombeck, and David W Tank. Intracellular dynamics of hippocampal place cells during virtual navigation. Nature, 461(7266):941–946, 2009.

40. Katie C Bittner, Christine Grienberger, Sachin P Vaidya, Aaron D Milstein, John J Macklin, Junghyup Suh, Susumu Tonegawa, and Jeffrey C Magee. Conjunctive input processing drives feature selectivity in hippocampal ca1 neurons. Nature neuroscience, 18(8):1133–1142, 2015.

41. Brad K Hulse, Evgueniy V Lubenov, and Athanassios G Siapas. Brain state dependence of hippocampal subthreshold activity in awake mice. Cell reports, 18(1):136–147, 2017.

42. Jeremy D Cohen, Mark Bolstad, and Albert K Lee. Experience-dependent shaping of hippocampal ca1 intracellular activity in novel and familiar environments. Elife, 6:e23040, 2017.

43. Kiryl D Piatkevich, Seth Bensussen, Hua-an Tseng, Sanaya N Shroff, Violeta Gisselle Lopez-Huerta, Demian Park, Erica E Jung, Or A Shemesh, Christoph Straub, Howard J Gritton, et al. Population imaging of neural activity in awake behaving mice. Nature, 574 (7778):413–417, 2019.

44. Xiaomin Zhang, Alois Schlögl, and Peter Jonas. Selective routing of spatial information flow from input to output in hippocampal granule cells. Neuron, 107(6):1212–1225, 2020.

45. Koichiro Kajikawa, Brad K Hulse, Athanassios G Siapas, and Evgueniy V Lubenov. Updown states and ripples differentially modulate membrane potential dynamics across dg, ca3, and ca1 in awake mice. Elife, 11:e69596, 2022.

46. S Wenceslao Evans, Dong-Qing Shi, Mariya Chavarha, Mark H Plitt, Jiannis Taxidis, Blake Madruga, Jiang Lan Fan, Fuu-Jiun Hwang, Siri C van Keulen, Carl-Mikael Suomivuori, et al. A positively tuned voltage indicator for extended electrical recordings in the brain. Nature methods, 20(7):1104–1113, 2023.

47. Jiannis Taxidis, Blake Madruga, Karen Safaryan, Conor C Dorian, Maxwell D Melin, Zoë Day, Michael Z Lin, and Peyman Golshani. Voltage imaging reveals hippocampal inhibitory dynamics shaping pyramidal memory-encoding sequences. Nature Neuroscience, 28(9): 1946–1958, 2025.

48. Jeffrey S Anderson, Ilan Lampl, Deda C Gillespie, and David Ferster. The contribution of noise to contrast invariance of orientation tuning in cat visual cortex. Science, 290(5498): 1968–1972, 2000.

49. Nicholas J Priebe and David Ferster. Direction selectivity of excitation and inhibition in simple cells of the cat primary visual cortex. Neuron, 45(1):133–145, 2005.

50. Hidehiko K Inagaki, Lorenzo Fontolan, Sandro Romani, and Karel Svoboda. Discrete attractor dynamics underlies persistent activity in the frontal cortex. Nature, 566(7743):212–217, 2019.

51. Paul K LaFosse, Zhishang Zhou, Jonathan F O’Rawe, Nina G Friedman, Victoria M Scott, Yanting Deng, and Mark H Histed. Cellular-resolution optogenetics reveals attenuation-by-suppression in visual cortical neurons. Proceedings of the National Academy of Sciences, 121(45):e2318837121, 2024.

52. Ido Aizenbud, Daniela Yoeli, David Beniaguev, Christiaan PJ de Kock, Michael London, and Idan Segev. Dendritic morphology and synaptic nonlinearities enhance functional complexity in human cortical neurons. Proceedings of the National Academy of Sciences, 123(28): e2533168123, 2026.

53. Michael E Hasselmo. The role of acetylcholine in learning and memory. Current opinion in neurobiology, 16(6):710–715, 2006.

54. Michael E Hasselmo, Clara Bodelón, and Bradley P Wyble. A proposed function for hippocampal theta rhythm: separate phases of encoding and retrieval enhance reversal of prior learning. Neural computation, 14(4):793–817, 2002.

55. Misha V Tsodyks, William E Skaggs, Terrence J Sejnowski, and Bruce L McNaughton. Paradoxical effects of external modulation of inhibitory interneurons. Journal of neuroscience, 17(11):4382–4388, 1997.

56. Laurel Watkins de Jong, Mohammadreza Mohagheghi Nejad, Euisik Yoon, Sen Cheng, and Kamran Diba. Optogenetics reveals paradoxical network stabilizations in hippocampal ca1 and ca3. Current Biology, 33(9):1689–1703, 2023.

57. Alessandro Sanzeni, Bradley Akitake, Hannah C Goldbach, Caitlin E Leedy, Nicolas Brunel, and Mark H Histed. Inhibition stabilization is a widespread property of cortical networks. Elife, 9:e54875, 2020.

58. Y Ahmadian, DB Rubin, and KD Miller. Analysis of the stabilized supralinear network. Neural Computation, 25(8):1994–2037, 2013.

59. DB Rubin, SD van Hooser, and KD Miller. The stabilized supralinear network: a unifying circuit motif underlying multi-input integration in sensory cortex. Neuron, 85(2):402–417, 2015.

60. Paul J. Goulart and Yuwen Chen. Clarabel: An interior-point solver for conic programs with quadratic objectives, 2024.

61. Hirotugu Akaike. A new look at the statistical model identification. IEEE transactions on automatic control, 19(6):716–723, 1974.

62. Gideon Schwarz. Estimating the dimension of a model. The annals of statistics, pages 461–464, 1978.

